# Ancestry adjustment improves genome-wide estimates of regional intolerance

**DOI:** 10.1101/2020.03.05.979203

**Authors:** Tristan J. Hayeck, Nicholas Stong, Evan Baugh, Ryan Dhindsa, Tychele N. Turner, Ayan Malakar, Timothy L. Mosbruger, Yuncheng Duan, Iuliana Ionita-Laza, David Goldstein, Andrew S. Allen

## Abstract

Genomic regions subject to purifying selection are more likely to carry disease causing mutations. Cross species conservation is often used to identify such regions but has limited resolution to detect selection on short evolutionary timescales such as that occurring in only one species. In contrast, intolerance looks for depletion of variation relative to expectation within a species, allowing species specific features to be identified. When estimating the intolerance of noncoding sequence methods strongly leverage variant frequency distributions. As the expected distributions depend on demography, if not properly controlled for, ancestral population source may obfuscate signals of selection. We demonstrate that properly incorporating demography in intolerance estimation greatly improved variant classification (13% increase in AUC relative to comparison constraint test, CDTS; and 9% relative to conservation). We provide a genome-wide intolerance map that is conditional on demographic history that is likely to be particularly valuable for variant prioritization.

## Introduction

Understanding the functional impact of noncoding sequence on protein coding sequence is one of the largest challenges in human genomics. Our ability to call variation in noncoding sequence has greatly outpaced our ability to interpret that variation and, currently, even studies employing whole genome sequencing (WGS) often restrict analyses to coding sequence. Previously, cross-species conservation has been used to identify genomic regions of likely importance. These methods are effective at identifying genomic regions that retain their functional importance across different species,^1–4^ but are not effective at identifying genomic regions that have emerged as important in a given species.^5^ However, emerging WGS datasets present an opportunity to address this problem as they provide a mechanism for detecting signatures of purifying selection within noncoding sequence by looking for intolerance in large standing human populations,^6,7^ where up until recently this has been difficult to detect due to relatively small sample sizes.

Methods for estimating genetic intolerance have previously been applied to noncoding sequence by either by comparing the observed local distribution of variation to expectation under neutrality given a sequence context informed mutation rate,^6^ or by comparing local sequence context dependent distributions of variation to genome-wide sequence context dependent distributions.^7^ One such method, Orion,^6^ was shown to be highly discriminative of known classes of regulatory elements and in a recent machine learning based classifier^8^, it was shown to be the most informative feature of variant pathogenicity among a set of features that characterize intolerance, conservation, 3D structure, expression, and other combined metrics. Here, we improve upon existing methods in several ways. First, instead of comparing the observed SFS to that expected under neutrality, we compute the expectation empirically, by stochastically sampling from putatively neutral regions making the method less sensitive to demographic factors that may distort the SFS, as these factors will affect both the observed SFS and its neutral expectation. Second, we stratify the analysis by ancestry, computing both the observed SFS and its expectation within each ancestry and then combining these contrasts into the final Population Conditional Intolerance Test (PCIT). It is well known that genetic diversity varies across ancestries and natural selection drives population differences in disease response.^9^ By stratifying the analysis on ancestry we effectively eliminate variability in genetic diversity between human subpopulations in estimating intolerance, leading to greater precision, as we demonstrate below.

## Results

### Differences in Neutral SFS Across Ancestry

We begin by investigating whether the neutral SFS varies across ancestry. Since our approach contrasts the observed SFS within a region to an empirical estimate of the neutral SFS, if there are differences in the neutral SFS across ancestry groups, stratifying on ancestry could eliminate an important source of variability leading to increased power. To this end, we used 15,708 whole genome sequenced samples from the genome aggregation database (gnomAD) across 8 different ancestries,^10^ including: African, American Latino/Admixed American, Ashkenazi Jewish, East Asian, Finnish, Non-Finnish European, South Asian, and other.

We estimate the neutral SFS by sampling variation from an intergenic sequence not annotated to be functional (see Quality Control, Defining Intergenic Regions, and Annotations). To test whether a given subpopulation’s neutral SFS differs significantly from another subpopulation we employ the following approach: begin by randomly sampled a million positions from neutral sequence and then randomly assigned a given position to be part of the SFS estimation for one of the two subpopulations being compared. This gives half a million positions on which to estimate the neutral SFS for each subpopulation, then quantify the difference between the two distributions using the log-rank test. This process was repeated a thousand times, the subpopulation label was then randomly shuffled at each site and the log rank test was recalculated after assignment. This gives us a null distribution of SFS differences between subpopulation. This was done with each pairwise ancestry, within each ancestry, and using the entire combined gnomAD population (Fig. 1c).

**Fig. 1.**
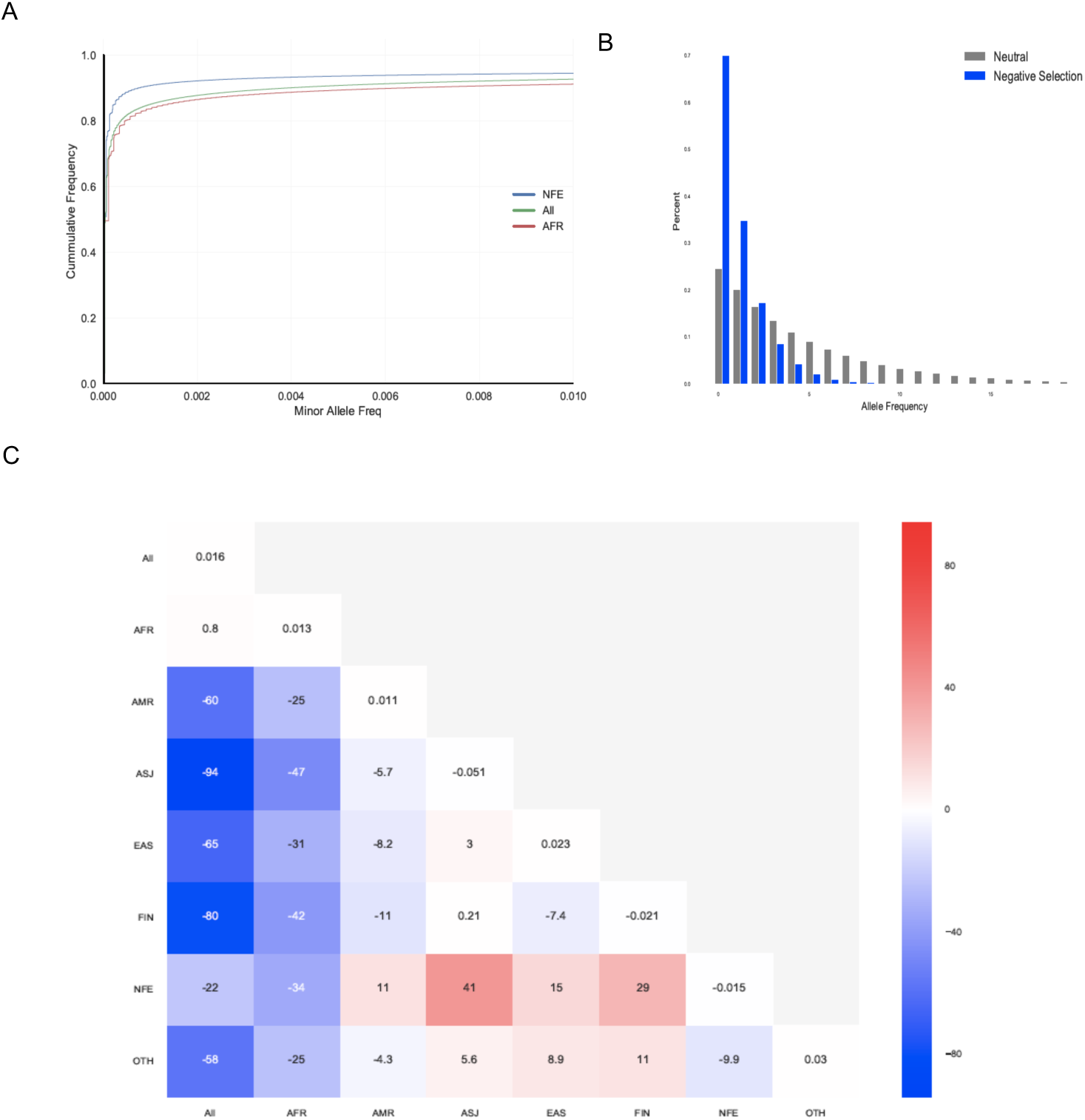
Characterizing cross population shifts in the site frequency spectrum. Understanding cross population shifts in site frequency spectrum, first is an illustrative diagram a) adapted from Sawyer and Hartl 1992 Genetics (remove Hartl version and put in ours) showing how negative selection is expected to effect the SFS. Then an empirical sample of b) the cumulative SFS for million intergenic bases taken across the two largest populations in gnomAD and the combined cumulative SFS. In the lower plot is c) a heatmap of the average over a thousand random permutations of half a million intergenic positions in one population versus another half a million intergenic positions from another population.

The African/African-American (AFR) SFS does not differ significantly from the pooled gnomAD SFS sample population (average logrank across thousand iterations: 0.8: p-value = 0.21), whereas every other population demonstrates a significant difference in SFS relative to the full population. It is to be suspected that the AFR population’s neutral SFS would demonstrate the most genetic diversity (Fig. 1b). When looking at the largest ancestries, AFR and Non-Finish European (NFE) samples, there is a significant difference in their SFSs (average logrank test - 34: p<1e-8). The differences observed between the neutral SFSs across gnomAD ancestry groups suggest that conditioning on ancestry in intolerance estimation may control an important source of variability, leading to improved precision. We investigate this approach in the next section.

### Ancestry Adjusted Intolerance Estimation

We conducted a genome wide scan of regional intolerance by looking for differences in the SFS within a given genomic region with an ancestry specific estimate of the neutral SFS. Specifically, we estimated the neutral SFS from a million random intergenic positions that were stochastically sampled across the genome, within a given ancestry and across all ancestries. We then compared this neutral SFS to the SFS distribution estimated from a given hundred and one base pair window (fifty bases on each side of the index position) using a weighted stratified log-rank test. The strata in the log-rank test are the different ancestries and are weighted by the inverse of the proportion of individuals from a given subpopulation in the gnomad (see online methods) to get the population conditional intolerance test (PCIT). Using a stratified test is a natural approach in settings like this where there are clear categorical groups, ancestral populations, that need to be controlled for to more accurately test for association. The motivation behind weighting the stratum is to reflect sampling where the various subpopulations are equally represented and to improve estimation of variance in the test statistic.^11,12^ We also conducted an unadjusted analysis where both the neutral SFS and the observed SFS within window were estimated from a pooled sample comprised of all ancestries. We refer to this as the unadjusted intolerance test (UIT). We then slide the hundred base query window to cover every position across the genome, with the restriction that the regions considered pass coverage and QC criteria (see Quality Control, Defining Intergenic Regions, and Annotations section). To improve computational efficiency, we take advantage of the fact that as you move from window to window very little of the data actually change and thus we can leverage our previous calculations in updating to a new window, avoiding the need to reload and recalculate across all data elements.

We investigated the performance of the various approaches by looking at how they classify non-coding ClinVar pathogenic variants versus a million randomly sampled common variants (MAF>5%) taken from 62,784 whole genome Trans-Omics for Precision Medicine (TOPMed projects) samples. As can be seen from figure 1, areas under the receiver operator curves (AUCs) are significantly improved when ancestry is accounted for (PCIT: AUC=91%) relative to when it is not (UIT: AUC=81%). PCIT also substantially outperforms two previously proposed approaches for measuring constraint and conservation in noncoding sequence (CDTS: AUC=78% and GERP: AUC=82%) (Fig. 2). Similar results are seen in classifying ClinVar coding variants. Specifically, we found with when ancestry is accounted for (PCIT: AUC=93%) relative to a similar unadjusted approach (UIT: AUC=85%). It also substantially outperforms a previously proposed approach for classifying noncoding sequence (CDTS: AUC=81% and GERP: AUC=90%) (Supp. Fig. 1). The same comparison was done with Orion, which also uses a sliding window approach to compare the observed SFS with a theoretical expectation under neutrality, computed given the sequence context dependent mutation rate found within the window. Orion slightly outperformed CDTS but underperformed relative to both the new population unadjusted intolerance test and the population conditional intolerance test (Orion non-coding AUC=78%, coding AUC =80%). Orion was run using a different quality control criteria and, as a result, used a significantly different set of variants. For this reason, we left it out of the comparisons found in the main text, but still note it in the supplementary materials (Supp. Fig. 2).

**Fig. 2.**
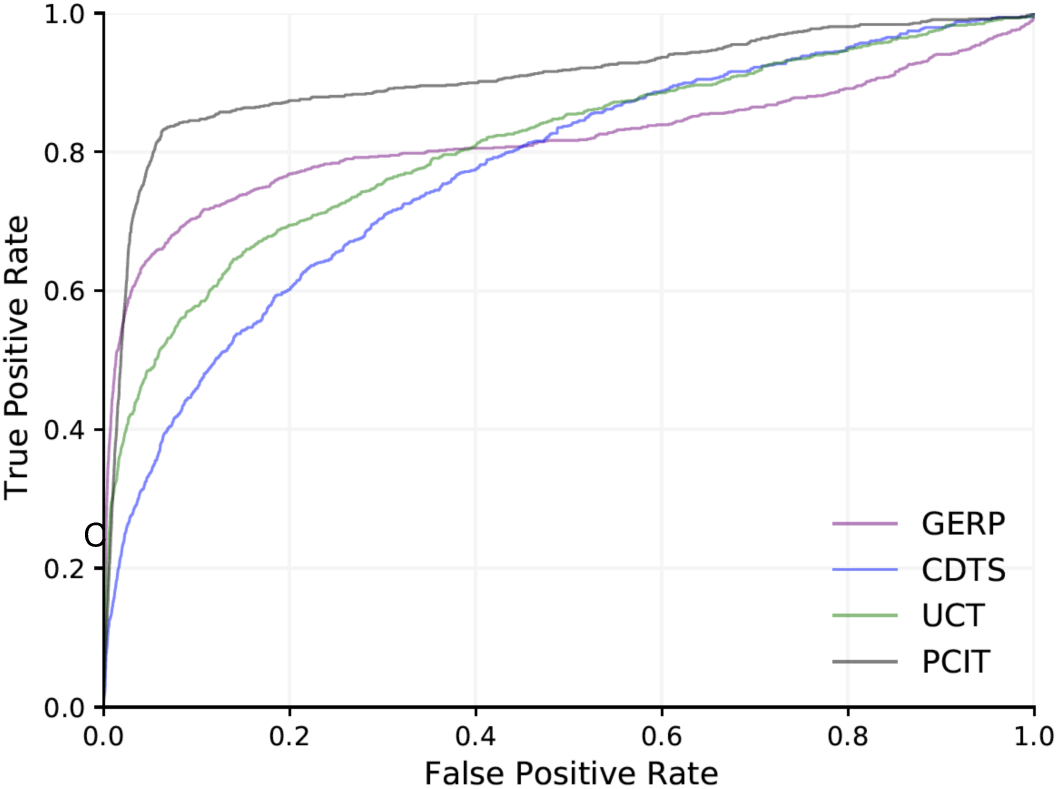
Predictive utility of population conditional constraint test relative to other constraint and conservation metrics for ClinVar non-coding variants. Non-coding ClinVar pathogenic variants versus a million randomly sampled variants from TOPMed with MAF greater than 5%.

We next investigated how sequences showing extreme PCIT scores (in the top 10%) are distributed across different genomic regions, such as exons: introns, enhancers, promotors, etc. by comparing how often sequence with a given annotation lies in the top decile of PCIT relative to sequence found in common intergenic regions (details of annotations described in Quality Control, Defining Intergenic Regions, and Annotations). We found that ultra-conserved regions have a 34.81 fold enrichment of extreme PCIT scores relative to common intergenic regions (Fig. 3). Exons show a 21.31 fold enrichment;24.40 for UTRs; 23.58 for introns; 23.25 for enhancers; and 22.93 for Hi-C experimental data that was taken from di Iulio used to validate CDTS^7^ (Supp Table 1). Non-coding RNA elements also showed an enrichment of intolerance relative to common intergenic regions (Fig. 3): miRNA showed a 23.07 fold enrichment; and a 22.41 fold enrichment for lincRNA (Fig. 3).

**Fig. 3.**
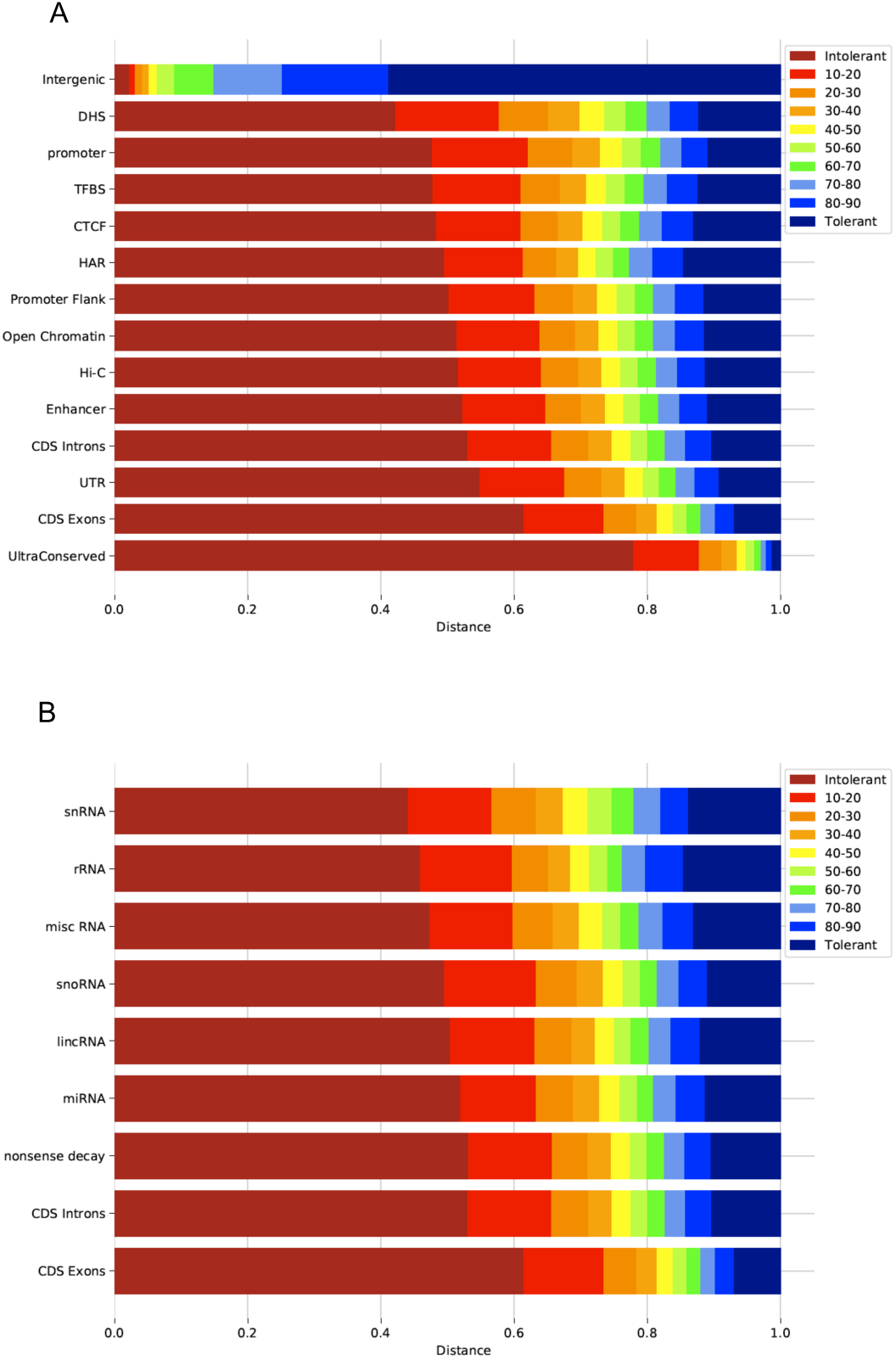
Relative constraint across different annotations from Ensembl^27^ and other resources.^5,6,31,32^. Decile break downs of PCIT scores across different functional annotations from and a million randomly sampled intergenic variants from TOPMed with MAF greater than 5%.

We then looked at how the intolerance of regulatory elements correlates with the genic intolerance of the genes they regulate. We began by connecting specific enhancer regions to the genes they regulate in various tissue types using Roadmap (http://www.biolchem.ucla.edu/labs/ernst/roadmaplinking/).^13–15^ As the PCIT is a nucleotide level score, we took the average of such scores across each enhancer, to create a single score per enhancer. Using the roadmap links for a given cell line, we linked a target gene for each enhancer. We binned the intolerance scores for enhancers targeting genes in a given gene set into 20 equal sized bins and took the median enhancer score within each bin. We also computed median RVIS score for the genes being targeted by the enhancers within each bin. Finally, we correlated the bins’ median enhancer and median target gene RVIS scores (Fig. 4). Consistent patterns of correlation between genic intolerance and regional regulatory intolerance were seen in OMIM genes (Fig. 4A-B correlation brain = 0.90, neurosphere 0.77), haploinsufficient genes (Fig. 4C-D correlation brain =0.65, neurosphere 0.51), and neurodevelopmental autosomal dominant genes (Fig. 4E-F correlation brain = 0.41, neurosphere 0.12).).

**Fig. 4.**
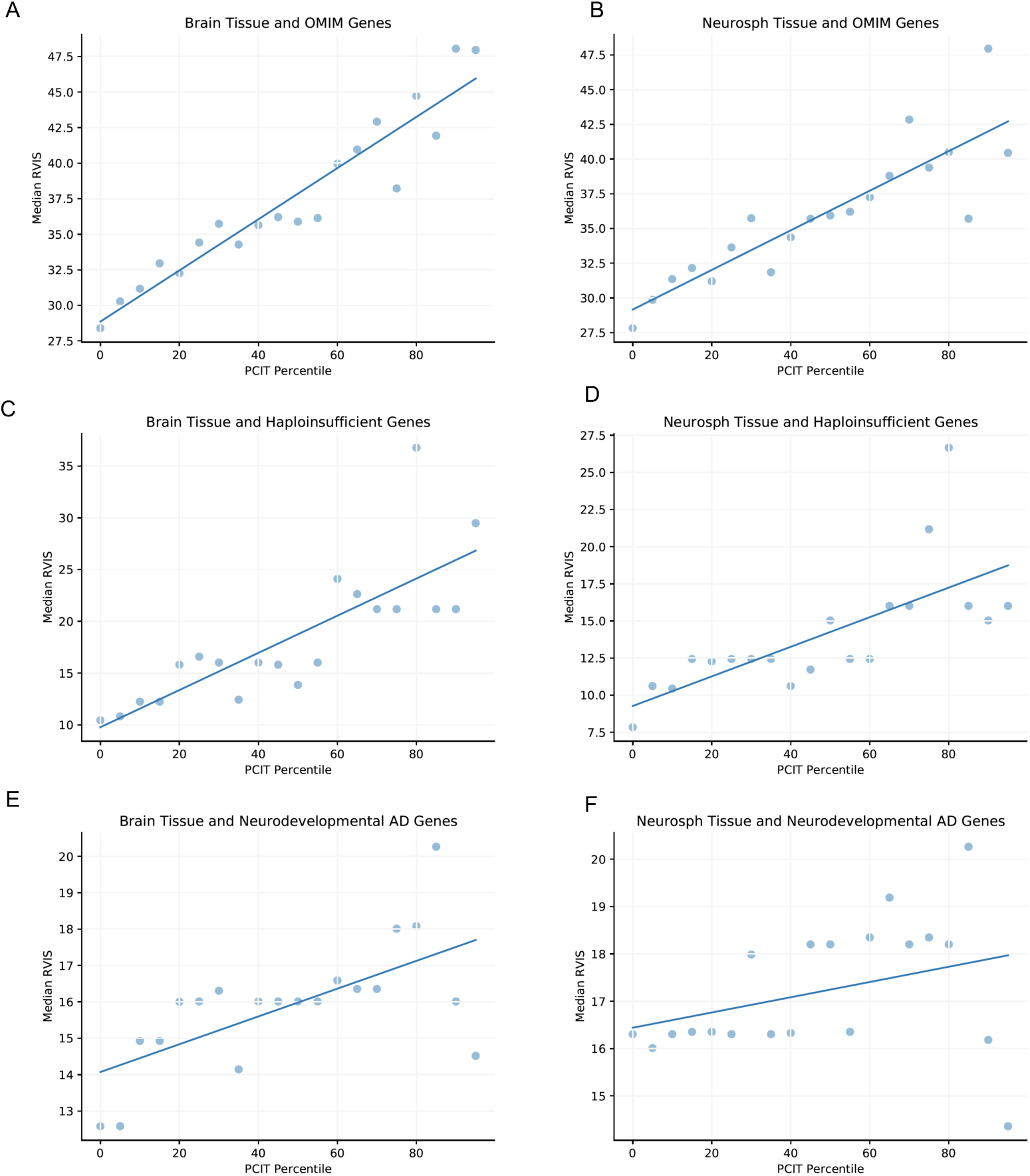
Genic intolerance to variation relative to PCIT constraint percentile of associated enhancers. Gene enhancer pairs were defined using Roadmap links,^12–14^ then the average SFS test constraint levels across enhancers where binned for every 5% then plotted versus the median RVIS scores. The top row corresponds to: (A-B) OMIM genes restricting to Roadmap A.) brain cells (Brain group) then B.) neurosphere cells, (C-D) haploinsufficient genes restricting to Roadmap C.) brain cells (Brain group) then D.) neurosphere cells, and finally (E-F) neurodevelopmental autosomal dominant genes restricting to Roadmap C.) brain cells (Brain group) then D.) neurosphere cells.

The strong correlations seen in the OMIM gene sets for brain and neurosphere linked enhancers were also seen when linked through other cell types with very strong correlations being seen when enhancers were linked via Epithelial (correlation =0.96) and muscle (muscle correlation = 0.93 and Myosat correlation = 0.90,) (Supp. Fig. 3).

## Discussion

Clear patterns of correlation were observed between intolerant regulatory elements and the genic intolerance scores of the target genes those elements regulate. The difference in more recent constraint captured using PCIT relative to conservation, as captured by GERP, appears to be larger in the context of non-coding variation as opposed to coding variation (Fig 2 and Supp. Fig. 1-2) which is consistent with recent findings that GERP may miss over half of non-coding mutations under purifying selection^17^. However, since there is limited known non-coding pathogenic variation more investigation needs to be performed.

Sample size is a key factor in being able to precisely identify intolerant regions. Given the current limited number of publicly available whole genome sequences, some ancestries are under-represented in this sample. For example, there are only 151 Ashkenazi Jewish individuals represented in the current analysis. However, with emerging population-based sequencing programs such as All of Us^18^ and the UK biobank^19^ we expect the number of sequences to increase dramatically in the coming years and to better represent diverse ancestries. This, in turn, will allow more precise regional intolerance estimation of smaller and smaller subunits of the genome. In the current study a sliding window of 100 bases was chosen to capture enough variation to precisely estimate intolerance while also balancing the localization of the scores. With larger and larger samples sizes, smaller windows will provide the same precision while better identifying finer and finer local structure in intolerance signal.

An alternative to the sliding window approach is to use predefined regional definitions based on known biology, e.g., enhancers, promotors, exons, introns, etc. However, the sizes of such regions can vary dramatically and, as a result, the precision of intolerance estimates will vary dramatically as well, with smaller regions’ intolerance often being very poorly estimated. A possible solution to this is to develop hierarchical models in this context that allow borrowing of information across similar regions, potentially stabilizing estimates. Such an approach has been successfully applied to intolerance estimation of subregions in coding sequence.^20^

Other methods for estimating regional intolerance consider sequence context variation rates, either to estimate an expected number of variants within a given region using a sequence context mutation rate,^10,21,22^ or by comparing a local rate of variation within a given sequence context to that observed in that context genome-wide.^7^ Though we found that our SFS-based approach outperformed CTDS, it is clear that two regions that are subject to the same level of purifying selection may have very different SFSs simply due to sequence context mutation rate differences between the regions. Thus, there is an opportunity to further refine the PCIT framework by developing sequence context informed SFS estimates within ancestry groups and to contrast that with an ancestry specific sequence context informed neutral SFS.

We do not directly compare to other methods that aggregate annotations^8,23–26^ from other rich feature sets^27–29^; however, we are encouraged that the approach proposed here will be useful in this context by the fact that Orion, a previously proposed SFS-based approach for estimating genome-wide regional intolerance, was recently demonstrated to be the most informative feature in a recent predictive model of variant pathogenicity.^8^ Improving such predictions, especially for variants in non-coding sequence, has implications for the interpretation of variation across genetics studies from genetic discovery to the diagnostic interpretation of patient genomes.

### Online Resources

https://github.com/tris-10/PopCondIntolTest

https://macarthurlab.org/2017/02/27/the-genome-aggregation-database-gnomad/

https://egg2.wustl.edu/roadmap/web_portal/chr_state_learning.html#core_15state

https://www.nhlbi.nih.gov/science/trans-omics-precision-medicine-topmed-program

http://www.biolchem.ucla.edu/labs/ernst/roadmaplinking/

### Quality Control, Defining Intergenic Regions, and Annotations

Variants included in SFS calculations for the PCIT had to meet gnomAD PASS criteria, the test was run on gnomAD v2.1.1. In addition, indels were excluded and only autosomal chromosomes were used. Low coverage regions where all positions in a window did not have 10X coverage in 70% of samples, SEGDUP regions, and recent repeat regions defined as having sequence < 10% diverged from the consensus in RepeatMasker^30^were removed. Aggarwala and Voight characterized sequence context and modeled heptamere mutation rates focusing on intergenic regions to explicitly avoid the potential impact of negative selection.^9^ We build on this definition to create an empirical sample of within ancestry intergenic SFS spectrums defined as the full set of genomic sequence filtering out centromeric, telomeric, repetitive regions, gene deserts of length greater than 2MB, sequence not present in the combined accessibility mask of 1000 genomes. Additionally, we restricted regions to be least 1KB away from any gene.

The annotations in Fig. 3 were predominantly taken from Ensembl^27^ including: CDS exon, CDS intron, CCCTC-binding factor (CTCF), promoter flanking regions, open chromatin regions, transcription factor binding sites (TFBS), enhancer, promoter, untranslated regions (UTR), transcript nonsense mediated decay, lincRNA, miRNA, snoRNA, miscRNA, rRNA, and snRNA. Additional annotations that were used included: Human accelerated regions (HAR),^31^ ultra conserved elements (UCE) (https://www.ultraconserved.org/),^32^ Hi-C experimental data,^7^ DNase I hypersensitive (DHS),^6^ and a million randomly sampled variants from TOPMed (https://www.nhlbi.nih.gov/science/trans-omics-precision-medicine-topmed-program)^33^ with MAF greater than 5%.Cell specific enhancers were defined based on Roadmap Core 15-state model (5 marks, 127 epigenomes https://egg2.wustl.edu/roadmap/web_portal/chr_state_learning.html#core_15state).

### Online Methods

We use a weighted and stratified log rank test to test for differences between the SFS within a given query window and the SFS estimated from intergenic neutral sequence across multiple ancestral populations.

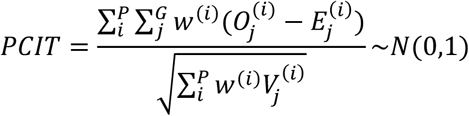

Using *P* different populations, separating the data into the different ancestral stratum, indexed by *i*, where within stratum we test the two groups (neutral and local window), G, using index j, where 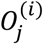 is the observed number of events, 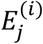 the expected number of events, and 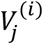 is the variance all using the ordinary log rank definitions.^34^ The failure events are taken to be the MAF for a given position and in this setting there is no right censoring. The weight, *w*^(*i*)^, was set to the inverse of the proportion of individuals from a given ancestry, where the weight was estimated by the number of individuals in the ancestral group relative to the total number of samples. The weighting was chosen looking at other schemes and verifying inverse proportions gave the best classification of pathogenic variation. To estimate the intergenic neutral SFS, we sampled a million random intergenic positions, and computed the SFS from the variation at those positions within each population. Our testing approach was optimized to take advantage of the fact that the data is sparse and does not change dramatically when moving from query window to query window. Thus, when moving from one window to another we can avoid fully reloading the data structures by simply removing the counts from the old position and then adding those for the new position. This greatly increases computational efficiency

