## Supplemental Materials for "Ancestry adjustment improves genome-wide estimates of regional intolerance"

**Supplement**

| *Annotation* | *Fold Enrichment* |
|---:|---:|
| *UltraConserved* | 34.64 |
| *CDS Exons* | 27.31 |
| *UTR* | 24.40 |
| *nonsense decay* | 23.64 |
| *CDS Introns* | 23.58 |
| *Enhancer* | 23.25 |
| *miRNA* | 23.07 |
| *Hi-C* | 22.93 |
| *Open Chromatin* | 22.86 |
| *lincRNA* | 22.41 |
| *Promoter Flanking* | 22.33 |
| *snoRNA* | 22.03 |
| *HAR* | 22.01 |
| *CTCF* | 21.50 |
| *TFBS* | 21.26 |
| *promoter* | 21.19 |
| *misc RNA* | 21.03 |
| *rRNA* | 20.44 |
| *snRNA* | 19.60 |
| *DHS* | 18.75 |

***Supp Table 1*** *relative enrichment of the across annotations of percent of scores in the top 10% in each annotation relative to common intergenic regions scores in the top 10%.*

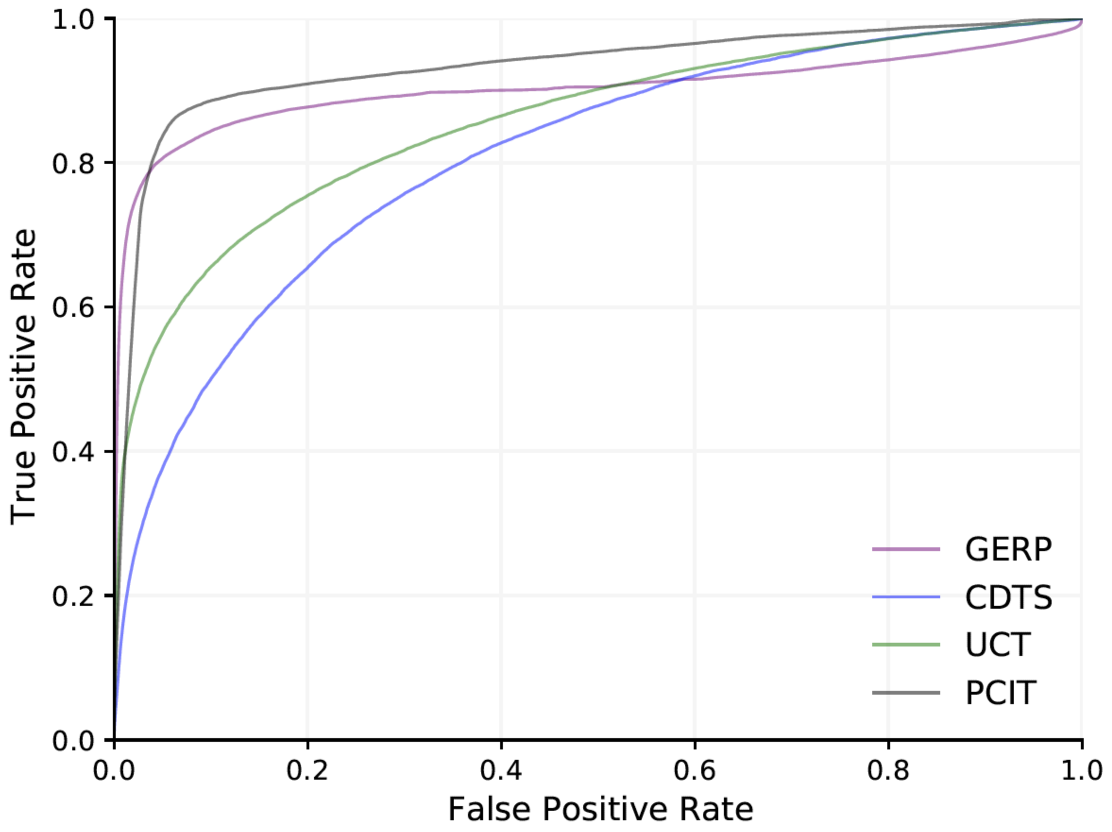


***Supp. Fig. 1*** ***Predictive utility of PCIT test relative to other constraint and conservation metrics for ClinVar coding variants.*** *Coding ClinVar pathogenic variants versus a million randomly sampled variants from TOPMed with MAF greater than 5%*

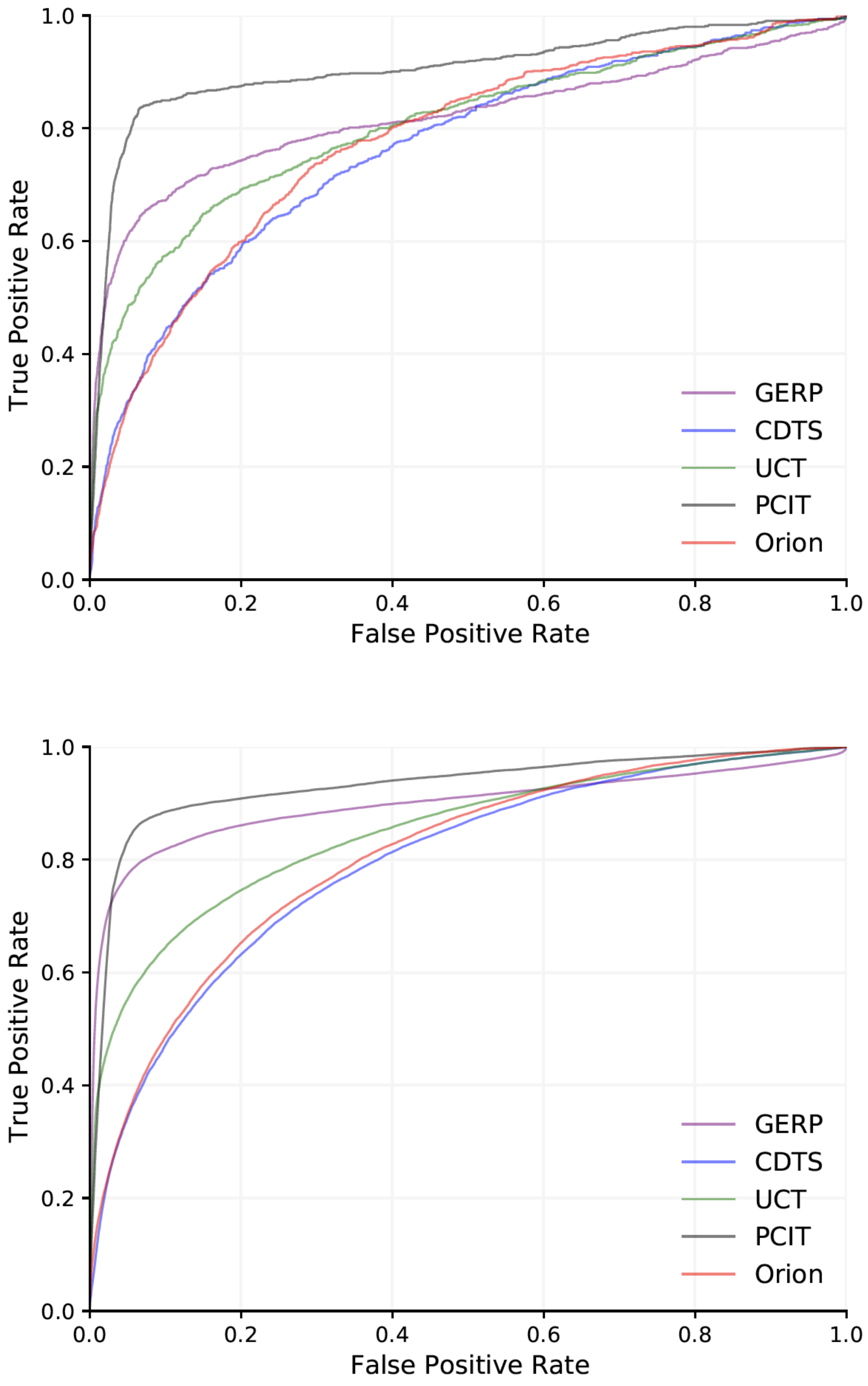


***Supp. Fig. 2* *Predictive utility of population conditional constraint test relative to other constraint and conservation metrics for ClinVar coding and non-coding variants restricting to non-repeat regions Orion is fit on.*** *ROC plot of different scores comparing A) Non-coding ClinVar pathogenic variants versus a half a million randomly sampled variants from TOPMed with MAF greater than 5% B) coding ClinVar pathogenic variants versus the same common variants.*

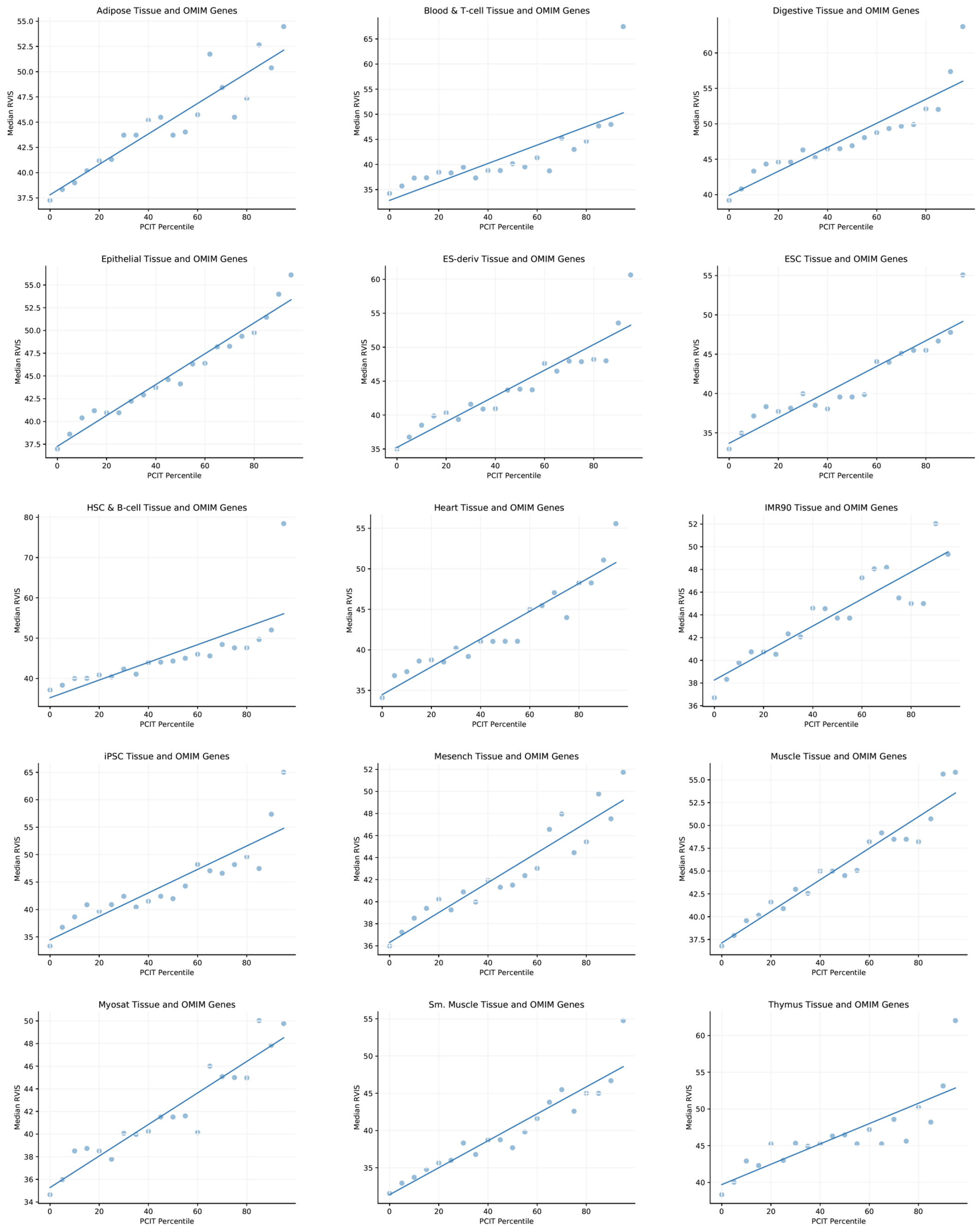


***Supp. Fig. 3 Genic intolerance in OMIM genes relative to PCIT constraint percentile of associated enhancers across multiple tissue types.** Gene enhancer pairs were defined using Roadmap links, then the average SFS test constraint levels across enhancers where binned for every 5% then plotted versus the median RVIS scores. Each tissue type is then restricted to OMIM genes.*
